# Computer Modeling of Radiofrequency Cardiac Ablation including Heartbeat-Induced Electrode Displacement

**DOI:** 10.1101/2021.12.11.472212

**Authors:** Juan J. Pérez, Ana González-Suárez, Enrique Nadal, Enrique Berjano

**Author notes:** **Corresponding author:** Enrique Berjano, Department of Electronic Engineering (Building 7F), Universitat Politècnica de València, Camino de Vera, 46022 Valencia, Spain.

## Abstract

**Background:** The state of the art in computer modeling of radiofrequency catheter ablation (RFCA) only considers a static model, i.e. it does not allow modeling ablation electrode displacements induced by tissue movement due to heartbeats. This feature is theoretically required, since heartbeat-induced changes in contact force can be detected during this clinical procedure.

**Methods:** We built a 2D RFCA model coupling electrical, thermal and mechanical problems and simulated a standard energy setting (25 W – 30 s). The mechanical interaction between the ablation electrode and tissue was dynamically modeled to reproduce heartbeat-induced changes in the electrode insertion depth from 0.86 to 2.05 mm, which corresponded with contact forces between 10 and 30 g when cardiac tissue was modeled by a hyperelastic Neo-Hookean model with a Young’s modulus of 75 kPa and Poisson’s ratio of 0.49.

**Results:** The dynamic model computed a lesion depth of 5.86 mm, which is within the range of previous experimental results based on a beating heart for a similar energy setting and contact force (5.6−6.7 mm). Lesion size was practically identical (differences less than 0.02 mm) to that using a static model with the electrode inserted to an average depth (1.46 mm, equivalent to 20 g contact force).

**Conclusions:** The RFCA dynamic model including heartbeat-induced electrode displacement predicts lesion depth reasonably well compared to previous experimental results based on a beating heart model, however this is true only at a standard energy setting and moderate contact force.

## 1. Introduction

Radiofrequency (RF) catheter ablation (RFCA) is intended to treat cardiac arrhythmias by applying RF electrical currents through an active electrode on the tip of a vascular catheter. Once the arrhythmia origin site has been located by electrophysiological mapping, RF electrical current flows between the active electrode and a dispersive electrode on the patient’s back, creating an irreversible thermal lesion in the target zone. Computer modeling has been shown to be a valuable tool for the study of electrical and thermal performance during RF ablation, not only to treat cardiac arrhythmias [1,2], but also for other purposes, such as tumor ablation [3]. Computational modeling of mechanical problems associated with tissue heating is currently a topic of interest [4−6].

RFCA computer models always consider that the active electrode is resting on the surface of the cardiac tissue and therefore remains slightly inserted. The insertion depth of the electrode in the myocardium during radiofrequency (RF) cardiac ablation (RFCA) is known to affect thermal lesion size. The deeper the electrode, the greater the power applied directly to the cardiac tissue and the less circulating blood. In fact, catheters able to measure contact force (CF) between electrode and tissue are now in regular use in clinical practice in an attempt to take insertion depth into account [7]. The contact area between electrode and tissue, i.e. the contour resulting from applying a mechanical force on the tissue is probably much more important than CF, and is also obviously related to surface deformation [8], since it determines how much power is targeted on the myocardium and how much ‘lost’ in the circulating blood.

Most computer models for RFCA to date assume the electrode to be inserted into the tissue and that the entire penetrating portion of the catheter makes contact with the tissue, i.e. the electrode is stuck (sharp insertion) without considering superficial deformation (e.g. [9−13]). To our knowledge, there are only five theoretical models on the study of RFCA that consider deformation of the endocardial surface (elastic insertion) [14−18]. In the model proposed by Cao *et al* [14] the deformation was not the result of solving the mechanical problem associated with the electrode/tissue contact force but was measured from the side view of tissue deformation using X-ray projection imaging and then incorporating the contour information into the computer model. From a procedural point of view they caused surface deformation by inserting the electrode into the tissue to a known depth. Unfortunately, they did not measure the contact force associated with different insertion depths. The model proposed by Petras *et al* [15] included the surface deformation from a specific value of contact force. The relation between applied force and surface deformation was obtained for any surface point using an approximation commonly employed in the mechanical indentation problem. This meant that the mechanical governing equations were not solved, and that the electrical-thermal problem was solved only after building a ‘static’ deformation profile. In contrast, both Yan *et al* [16] and Singh and Melnik [17] did solve the mechanical problem associated with tissue deformation by considering the cardiac tissue as a hyperelastic material and using the Mooney–Rivlin model to characterize its stress–strain curve. Both studies modeled constant voltage ablation, unlike the currently used constant power protocol. Ahn and Kim [18] also used a hyperelastic model to analyze cardiac tissue deformation due to the catheter contact force. These mechanical models are really inspired by a static image of the electrode at a given time. In other words, despite the fact that they all have represented an advance in RFCA modeling, none has considered the dynamic behavior of the mechanical interaction, i.e. heartbeat-induced electrode displacement. Changes in the CF associated with systole-diastole heart movements are normally found during RFCA [19], and in fact in some *ex vivo* models electrode displacement has been mimicked by placing tissue samples on motorized motion-controlled platforms [20,21]. Our goal was to build an RFCA computer model including heartbeat–induced electrode displacement and to compare the computer results with those obtained from previous experiments based on a beating heart in similar conditions of energy and contact force. To our knowledge, this is the first computer model that includes this realistic feature.

## 2. Methods

### 2.1. Model geometry

We used a limited-domain 2D model as described in Irastorza *et al* [22]. Figure 1A shows the geometry of the model which consisted of a vertical electrode (7 Fr, 4 mm) on a fragment of cardiac tissue surrounded by blood. The heartbeat-induced electrode displacement was assumed to be an up-and-down vertical movement that changed insertion depth (ID) from 0.86 to 2.05 mm (measured from the tissue surface). This movement followed a sine-like time evolution with a 1 Hz frequency, which is equivalent to heart rate of 60 beats per minute. This ID range was that of a contact force ranging from 10 to 30 g measured at the plastic section of the catheter, which can be considered as moderate in clinical practice [23]. This dynamic range of ∼20 g is also observed in clinical practice due to the heartbeat (e.g. see Fig. 1 of [19]). The results from this dynamic model were compared to those obtained with a static model in which the electrode remained at rest at a depth of 1.45 mm (equivalent to 20 g), which is the average position.

**Figure 1.**
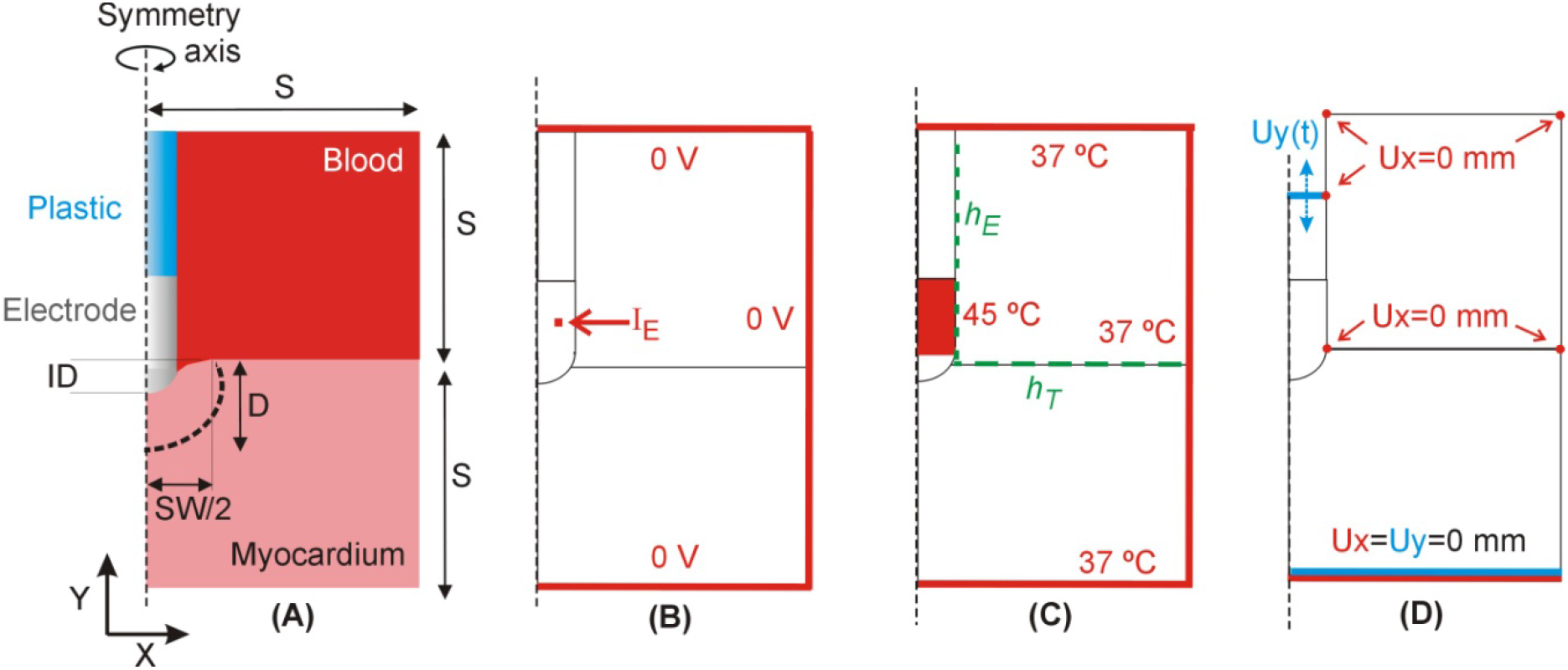
**A:** Geometry of the two-dimensional computational model built (not to scale) including an ablation electrode (7Fr, 4 mm) inserted into a fragment of myocardium and completely surrounded by blood. Dimension of myocardium and blood (S) is obtained from a convergence test. **B:** Electrical boundary conditions. The axial symmetry implies that the x-component of the current density is zero on the axis. **C:** Thermal boundary conditions. *h*_*E*_ and *h*_*T*_ are the thermal convection coefficients at the electrode–blood and tissue–blood interfaces, respectively. The axial symmetry implies that the x-component of the heat flow is zero on the axis. **D:** Mechanical boundary conditions of displacement (x-component in red and y-component in blue).

### 2.2. Governing equations

The computer model was based on a triple coupled electric-thermal-mechanical problem which was solved numerically using the Finite Element Method (FEM) with ANSYS software (ANSYS, Canonsburg, PA, USA). The governing equation for the thermal problem was the Bioheat Equation [24]:

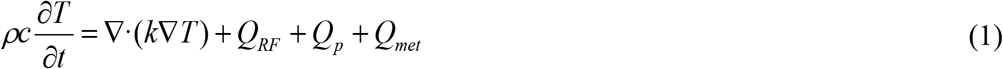

where *ρ* is density (kg/m^3^), *c* specific heat (J/kg·K), *T* temperature (ºC), *t* time (s), *k* thermal conductivity (W/m·K), *Q*_*RF*_ the heat source caused by RF power (W/m^3^), *Q*_*p*_ the heat loss caused by blood perfusion (W/m^3^) and *Q*_*m*_ the metabolic heat generation (W/m^3^). Both *Q*_*m*_ and *Q*_*p*_ were ignored as these terms are negligible compared to the others [24]. A quasi-static approximation was employed for the electrical problem. The magnitude of the vector electric field ***E*** was obtained from ***E*** = −∇Φ (Φ being voltage) while voltage was obtained from ∇·(*σ*(*T*)∇Φ) = 0 (*σ* being electrical conductivity). The distributed heat source *q* was then obtained as *Q*_*RF*_ = *σ*|***E***|^2^.

In order to model the vaporization in the myocardium, Eq. (1) was written as a balance of enthalpy changes instead of the energy changes proposed in [25]:

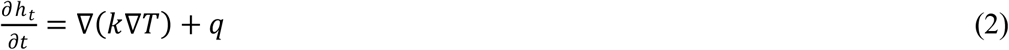

where *h*_*t*_ is the tissue enthalpy per unit volume. This value can be determined by assessing the amount of energy deposited in the tissue when its temperature is raised from 37 °C to values above 100 °C. According to [25], enthalpy per unit volume is:

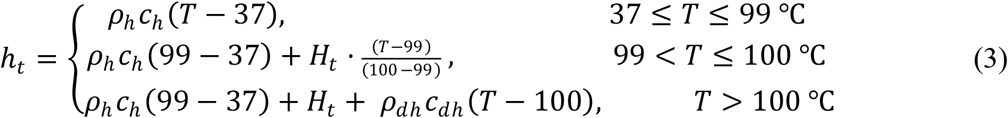

where the subscript *h* refers to the properties of the hydrated tissue (i.e. before reaching 99 ºC), the subscript *dh* refers to those of the dehydrated tissue, and *H*_*t*_ is the tissue vaporization latent heat. The partial derivative of the enthalpy in Eq. (2) can be therefore expressed as:

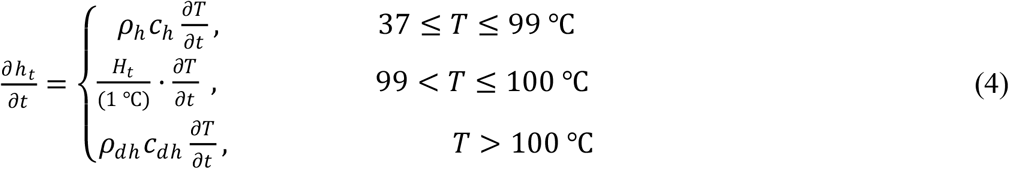

A complete mechanical model considering the electrode displacement caused by the myocardium movement should take the contact forces between the electrode tip and the myocardium into account. The contact forces depend on the mechanical material behavior, which depends on the material properties and on the inertial effects when dynamic effects are considered. We considered a hyperelastic mechanical model for the electrode and tissue, which could reproduce heartbeat-induced oscillatory electrode displacements. We were thus able to study the thermo-electrical effects due to the time variation of the contact area and the relative position of the electrode with respect to the tissue. Our model was quasi-static, i.e. viscosity and inertial effects were ignored. This is an approximation, since dynamic effects could appear. In mathematical terms, we modeled the large deformation elasticity problem considering the Lagrangian formulation. The solution of the elasticity problem follows by obtaining a displacement field compatible with the imposed displacements, such that the virtual work of the internal forces is the same as the virtual work of the external forces *δW*_*int*_ = *δW*_*ext*_, i.e. forces applied to the body. The virtual work of the internal forces is evaluated as follows:

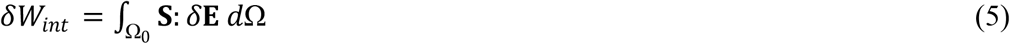

where **S** is the second Piola-Kirchoff stress tensor and *δ***E** is the Green’s tensor associated to the virtual displacement. The Piola-Kirchoff stress tensor can be written as follows

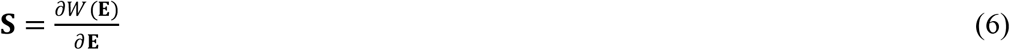

where *W* is the strain energy density function, which depends on the material considered for each solid. The metal (platinum) and polymeric components (polyurethane) of the catheter were considered as elastic solids with a Krichoff-Saint Venant material model, since their strains are small but they experience considerable displacement. The strain energy density function for this material is stated in the following equation:

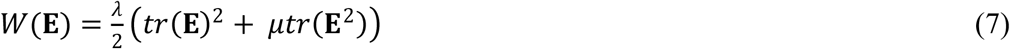

where *λ* and *μ* are the Lame’s constants that can be obtained through the following formulas:

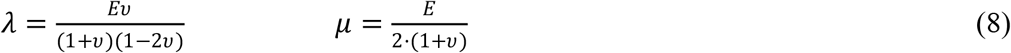

where *v* is the Poisson’s ration and *E* is the Young’s modulus. The Young’s modulus and Poisson’s ratio were 171 MPa and 0.39 for the metal electrode (platinum), respectively, and 10 MPa and 0.4 for catheter (polyurethane), respectively [26]. For the myocardium a hyperelastic Neo-Hookean model was considered with Young’s modulus of 75 kPa and Poisson’s ratio of 0.49 [15]. The strain energy density function was

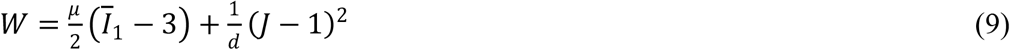

where the material properties are represented by *G*, the shear modulus, and *d=2/K* where *K* is the bulk modulus, *Ī*_1_ is the first invariant of the Cauchy-Green’s strain tensor (**C** = 2**E** + **I**) and *J* is the determinant of the strain gradient. Additionally, the relation between the bulk and the Young’s modulus and the Poisson’s ratio of a material can be obtained with the following expression:

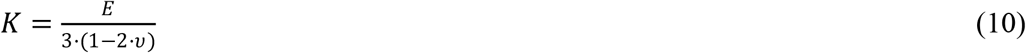

As a first approximation, the blood around the electrode and myocardium was assumed not to have any mechanical behavior. To simulate the movement of blood around the electrode and myocardium, a Neo-Hookean hyperelastic model was considered only for consistency in the simulation. For this, we set non-realistic mechanical material properties for the blood (Young’s modulus of 2 kPa and Poisson’s ratio of 0.4) with the only objective of avoiding numerical instabilities of the simulation while preserving no mechanical influence in the results. Several mechanical simulations were carried out to calibrate the model, comparing the results of the mechanical response (i.e. force applied to the electrode and displacements) with and without blood.

### 2.3. Electrical and thermal properties

The thermal properties of the plastic and metal elements of the catheter were taken from [27]. The thermal and electrical properties of the model elements are shown in Table 1 [27,28]. The electrical conductivity (*σ*) of myocardium was considered as a temperature-dependent function as follows: it rose exponentially +1.5%/ºC from 37 to 99 ºC (0.281 S/m at 37 ºC), then decreased two orders of magnitude from 99 to 100 ºC in order to model the drastic water loss due to vaporization (we previously demonstrated that using +1.5 or +2%/ºC, or 2 or 4 orders of magnitude produces very similar results [29]), and remained constant from 100 °C onwards. The thermal conductivity (*k*) of cardiac tissue was considered to be constant until reaching 99 ºC (0.56 W/K·m), and changed once the tissue was dehydrated (see Appendix [30−33]). The parameters used to model the phase change (vaporization) were as follows: *H*_*t*_ was estimated as the product of the water vaporization latent heat (*H*_*w*_) and the water mass fraction in the cardiac tissue (*C*). *H*_*w*_ was calculated as the product of the water vaporization latent heat (2256 kJ/kg) and water density (958 kg/m^3^) (both assessed at 100 ºC [31], given a value of 2.161×10^9^ J/m^3^. *C* was considered to be 75%, which is a typical value reported for heart and muscle [34,35]). *H*_*t*_ was therefore 1.62·10^9^ J/m^3^.

**Table 1.** Thermal and electrical characteristics of the elements employed in the model.

| Element/Material | $\sigma$ (S/m) | $k$ (W/m·K) | $\rho$ (kg/m<sup>3</sup>) | $c$ (J/kg·K) |
| --- | --- | --- | --- | --- |
| Electrode/Pt-Ir [27] | $4.6 \times 10^6$ | 71 | 21500 | 132 |
| Catheter/Polyurethane [27] | $10^{-5}$ | 23 | 1440 | 1050 |
| Cardiac Chamber/Blood [28] | 0.748 | 0.52 | 1050 | 3617 |
| Myocardium (hydrated [28]) | 0.281 | 0.56 | 1081 | 3686 |
| Myocardium (dehydrated)* | 0.007 | 0.287 | 1428 | 2474 |

$\sigma$ , electrical conductivity (at 500 kHz); $k$ , thermal conductivity; $\rho$ , density; and $c$ , specific heat (all assessed at 37 °C in case of tissue and blood). \* Estimated (see text and Appendix for details).

### 2.4. Initial and boundary conditions

The initial temperature in the entire model was 37 ºC. Fig. 1B and 1C show the electrical and thermal boundary conditions, respectively. The electrical current I_E_ injected at a node of the active electrode was adjusted at each time step in order to apply a constant RF power of 25 W for 30 s (note that the power value used in the simulations was reduced by 20% since the model did not include the entire torso, i.e. 20 W [22]). All the outer surfaces of the model were set to 0 V (Dirichlet boundary condition) except the surface of the symmetry axis, which was fixed at zero electric flux (Neumann boundary condition).

For the thermal boundary conditions, a null thermal flux was used on the symmetry axis and a constant temperature of 37 ºC was fixed on the outer surfaces of the model at a distance from the ablating electrode (this was also the initial temperature value). The effect of blood circulating inside the cardiac chamber was modeled by thermal convection coefficients at the electrode–blood (*h*_*E*_) and the tissue–blood (*h*_*T*_) interfaces, considering electrical conductivity of blood independently of temperature (as in Method 2 in [11]). The coefficients were calculated as in [12] for a blood flow of 0.085 m/s, simulating ablation sites with low local blood flow, as in patients with chronic atrial fibrillation and dilated atria [36]: *h*_*E*_ = 3346 W/m^2^·K and *h*_*T*_ = 610 W/m^2^·K. Electrode irrigation was modeled using a ‘reduced model’ as described in [13], which avoids solving the fluid dynamics problem by setting a constant temperature at the electrode tip. This approach is suitable for predicting lesion depth and maximum tissue temperature, but tends to overestimate surface width by ∼2−3 mm [13]. The irrigation of the electrode itself was modeled by fixing a temperature at 45 ºC only in the cylindrical zone of the electrode tip, leaving the semispherical tip free. This approximation for modeling an irrigated electrode is suitable for predicting lesion depth and the maximum temperature reached in the tissue at all times during ablation [13].

Fig. 1D shows the boundary conditions for the mechanical problem, specifically the displacement values in both axes (Ux and Uy) at relevant key points. We chose to use electrode displacement as load instead of CF for simplicity. Note that since the viscous behavior was not included, there is a direct and one-to-one relationship between CF and electrode vertical displacement. To state this relationship we first solved the static mechanical model for the three contact force values (10, 20 and 30 g) and obtained their corresponding displacement values on ANSYS, using a structural model and steady-state analysis (which means inertial forces and damping forces are ignored). The resulting electrode displacement values were used as peak values of the sinusoidal-waveform dynamic load applied on the catheter for the transient analysis. Figure 2 shows the consecutive time phases considered in each transient simulation (63 s in total): 1-s landing (the electrode comes into contact with the tissue surface), 30-s ablation (the electrode deforms the tissue surface by entering between 0.86 mm and 2.05 mm in the dynamic case –equivalent to a range of between 10 and 30 g of contact force) or remains steady at a depth of 0.86, 1.46 or 2.05 mm in the static cases (with contact forces of 10, 20 and 30 grams, respectively), 1-s takeoff (the electrode separates from the tissue), 30-s post-ablation (RF-off, in this period the lesion grows due to thermal latency), and 1-s landing (the electrode comes back into contact with the tissue surface). This last phase was simply conducted to measure the lesion size at the end of post-ablation period with the electrode inserted in the tissue.

**Figure 2.**
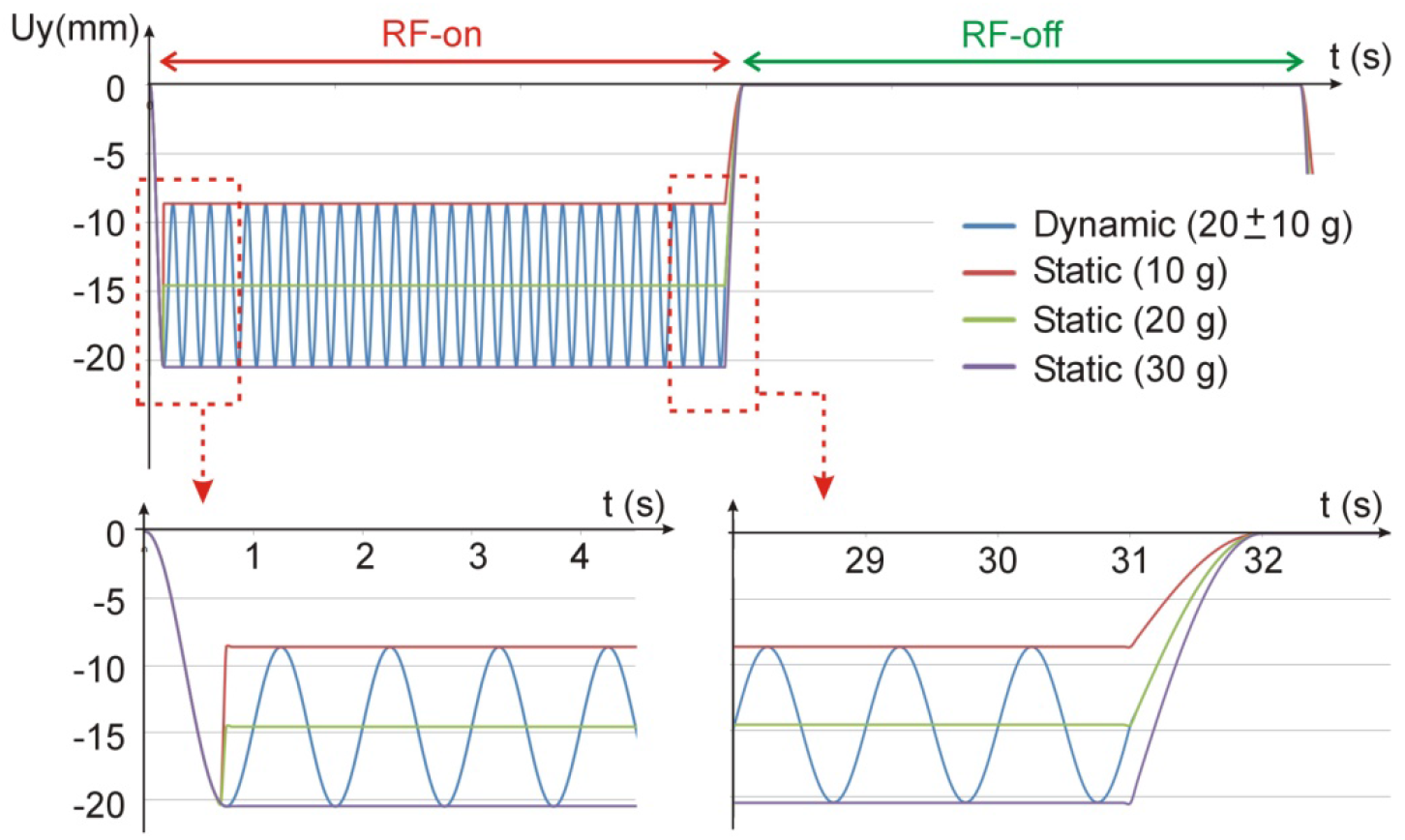
Evolution of the y-component of the displacement applied as a mechanical load on the catheter during the ablation time (RF-on) and post-ablation time (RF-off) for static and dynamic case and different values of contact forces.

The mechanical model was completed by considering the specific contact conditions between the three materials (electrode, myocardium and blood). The contact between electrode and myocardium considered the standard formulation with Signorini’s boundary conditions (i.e. avoiding penetration between surfaces, computing the surface pressure that guarantees no penetration condition, and allowing free separation). The contact between blood and myocardium followed a non-separation rule based on avoiding penetration between the materials, computing the surface pressure that guarantees no penetration condition, and once contact was established separation was not possible, thus normal tractions could be considered. Finally, a non-frictional contact model was used allowing free slip behavior between any pair of materials.

### 2.5. Model verification

Cardiac tissue and blood chamber dimensions (S in Fig. 1A) were previously estimated by means of a convergence test to avoid boundary effects and a value of S = 4 cm was found appropriate [22] using the value of the lesion depth (D) after 30 s of RFCA as a control parameter. Discretization was spatially heterogeneous: the finest zone was always the electrode-tissue interface with the largest voltage gradient and hence the maximum value of current density. The grid size was gradually increased in the tissue with distance from the electrode-tissue interface. We first considered a tentative spatial (i.e. minimum meshing size) and temporal resolution and determined the appropriate spatial resolution by means of a similar convergence test using the same control parameter as in the previous test. The mesh size was assumed to be suitable when an asymptotic tendency was seen in the lesion depth as mesh size decreased.

### 2.6. Output variables

The Arrhenius damage model was used to estimate lesion size from the temperature evolution computed at any point. This model associates temperature with exposure time by means of a first-order kinetics relationship:

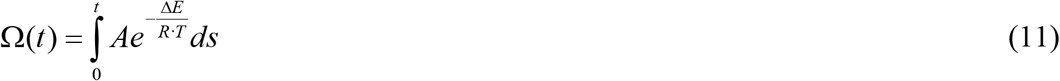

where *R* is the universal gas constant (8.314 J/K mol), *A* (7.39×10^39^ s^-1^) is a frequency factor and *ΔE* (2.577×10^5^ J/mol) is the activation energy for the irreversible damage reaction [27]. Lesion contour was estimated using the Ω = 1 isoline.

A ‘footprint’ is usually observed after ablation in both the thigh muscle model [36] and the beating heart model [23], possibly due to myocardium viscosity and to the stiffer tissue after denaturing. Since our model did not consider either of these two characteristics, the only way to mimic the footprint was to measure lesion size while the electrode was inserted, i.e. we measured the lesion depth (since the tissue surface, i.e. D in Fig. 1A) and surface width (SW in Fig. 1A) after the post-RF period at two almost simultaneous times: at 62 s when the electrode remained withdrawn (0 g contact force) and 1 s later (i.e. t=63 s) when the electrode was once more inserted to the same depth as during ablation (1.46 mm in the dynamic case).

## 3. Results

### 3.1. Verification of the model

The model had 7.791 nodes and 3,044 triangular elements. When the outer dimension (parameter S in Fig. 1A) was enlarged to 8 cm, the number of nodes and elements slightly varied to 8,709 and 3,422, respectively, but the change in lesion depth stayed below 0.22 mm and the maximum tissue temperature varied less than 1 ºC. Likewise, when the grid size was drastically reduced (around 30,000 nodes and more than 12,000 elements), lesion depth varied less than 0.02 mm and the maximum tissue temperature less than 0.5 ºC. The former values of mesh size and outer dimensions were thus considered to be suitable.

### 3.2. Comparison between static and dynamic contact force

Table 2 shows the depth (D) and surface width (SW) of lesions computed for the static and dynamic cases. There is hardly any difference between the measurements computed with the electrode inserted and withdrawn (see Figure 3): less than 0.33 mm for depth and 0.04 mm for surface width (these values can be considered insignificant since they are less <0.5 mm, which is approximately that of the deviation observed in experimental RFCA studies [11]. The lesion size obtained from the 20±10 g dynamic model is practically identical to that of the 20 g static model (differences less than 0.04 mm). Lesion size increased slightly when contact force was raised from 10 to 30 g: depth increased from 5.30 to 6.37 mm, and surface width increased from 4.38 to 4.52 mm (measurements with inserted electrode).

**Table 2.**
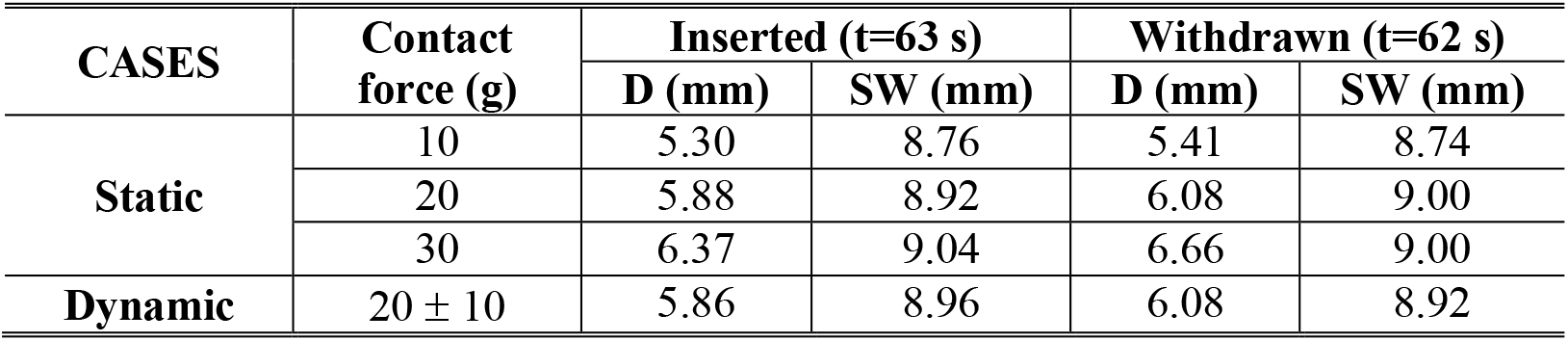
Depth (D) and surface width (SW) of lesions computed with the electrode inserted in the tissue and withdrawn for static and dynamic cases (20 W, 30 s).

**Figure 3.**
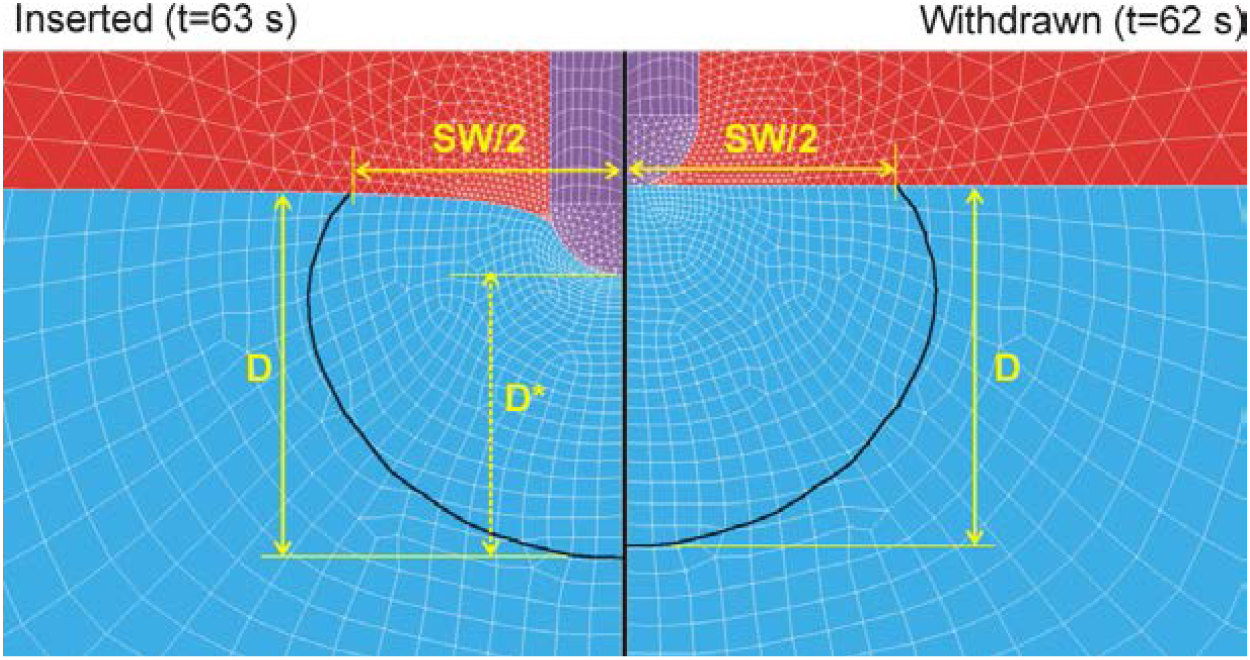
Thermal lesion contour computed with Arrhenius damage model (Ω = 1) for electrode inserted (A) and withdrawn (B) after the post-RF period. SW: surface width, D: depth from tissue surface, and D*: depth from electrode tip.

The dynamic performance of the temperature distributions in the myocardium is given in the video file in which the first 1-s landing phase is not shown since it was due to the mechanical bond between model elements: http://personales.upv.es/eberjano/Supplementary-material-Video.mp4 Figure 4 shows the temperature distributions at different times during ablation (up to 30 s) and post-ablation for the dynamic case with CF of 20±10 g. As can be seen in Fig. 4 and the video file, as the electrode periodically pushes the tissue, the nodes move with this deformation so that the entire temperature distribution oscillates slightly. The evolution of the temperature at a specific depth from the tissue surface will also have small superimposed oscillations.

**Figure 4.**
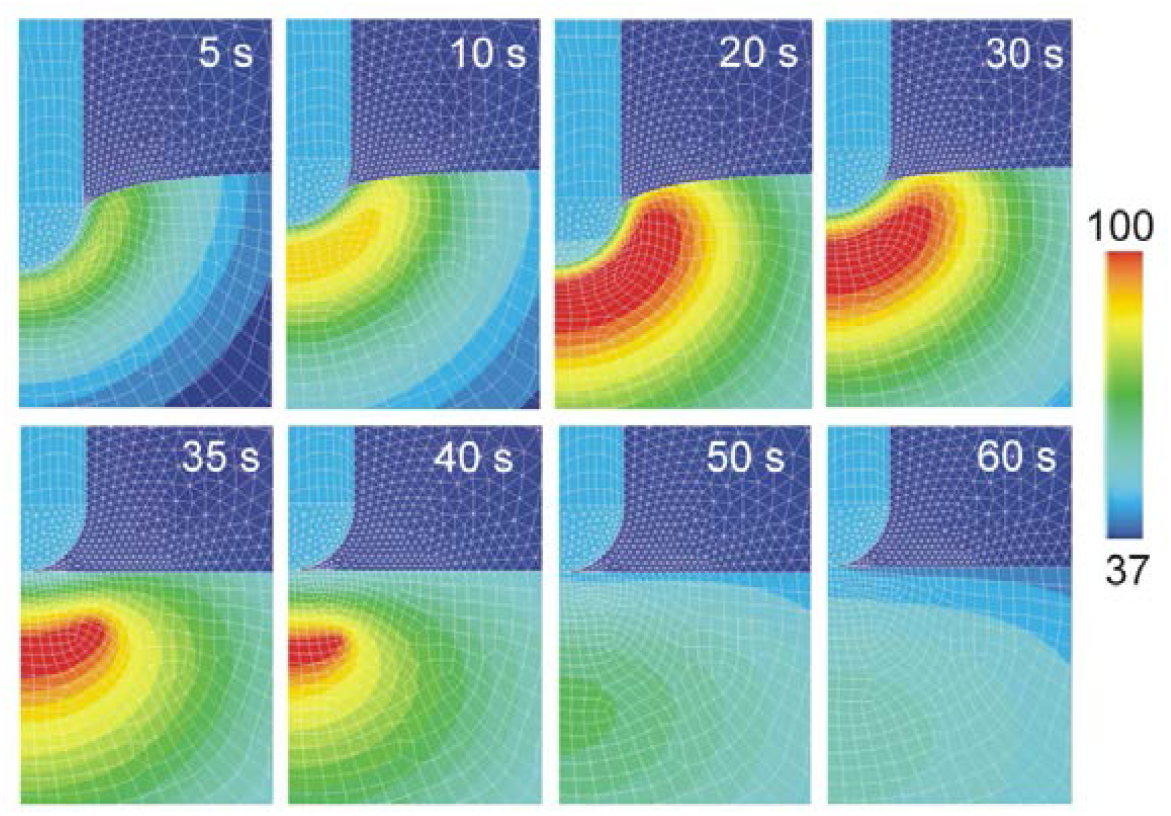
Temperature distributions in the tissue at different ablation times (5, 10, 20 and 30 s) and post-ablation period (35, 40, 50 and 60 s). The plots are those of different electrode positions within its vertical displacement in the dynamic case with a CF of 20±10 g. Scale in ºC.

### 3.3. Effect of modeling hydrated tissue

The model considered different values of thermal conductivity (*k*) and specific heat (*c*) for the myocardium before and after dehydration (see Appendix and Table 1). We repeated the computer simulations for the dynamic case keeping the same values of *k* and *c* above and below 100 ºC (i.e. 0.56 W/m·K and 3686 J/kg·K, respectively), obtaining differences in lesion size below 0.03 mm (note that *c* always increased between 99 and 100 ºC to model the latent heat associated with the phase change).

## 4. Discussion

### 4.1. Effect of heartbeat-induced electrode displacement

The aim was to build an RFCA computer model including heartbeat-induced electrode displacement. Until now, none of the models including mechanical tissue deformation considered the dynamic changes associated with the heartbeat [15−18]. While these changes can be noted in clinical practice and appear as variations in the contact force measured by the sensor at the tip of the catheter [19], our simulation results suggest that they have a limited impact on lesion size, although the temperature map in the tissue oscillates slightly so that the temperature at a given depth will also experience oscillations. We previously observed similar behavior in another simulation context in a study in which the applied RF voltage was oscillating at a low frequency (around 1 Hz) [37]. There we found that even though the temperature at a given depth oscillated with the oscillating RF voltage, the resulting lesion size was practically the same as when the oscillations were not considered. Our results thus suggest that in the specific case of moderate contact force (<30 g) and a standard energy setting (e.g. 25 W for 30 s) the dynamic and static model compute similar lesion size. This could be an advantage in terms of computational cost, since a model including the electrode displacement takes ∼16 hours of simulation (the results file occupies 6 GB) while a static model only takes 30 min (the results file occupies 155 MB).

### 4.2. Comparison with experimental data

Table 3 compares lesion size computed with the dynamic model and previous experimental results based on a beating heart model. The experimental data were chosen from those with similar contact force, power and duration [23,36,38]. The lesion depth computed by the dynamic model is within the range of the experimental results, while the lesion surface width agrees with the results of Leshem *et al* [38] and is ∼2 mm wider than those reported by Ikeda *et al* [23] and Nakawaga *et al* [36]. In this regard it is important to note that the ‘reduced model’ used in this study tends to overestimate the surface width by ∼2−3 mm [13].

**Table 3.**
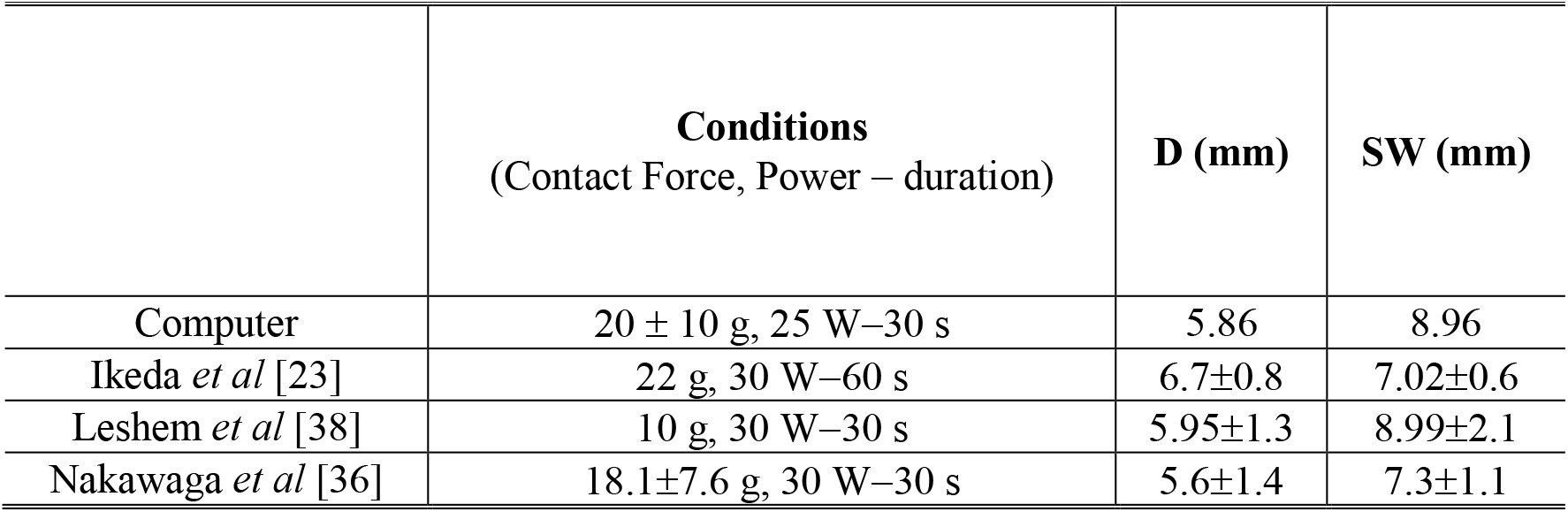
Comparison of computer (dynamic case) and experimental (beating heart model) results in terms of lesion size (D: depth, SW: Surface width).

|  | Conditions<br>(Contact Force, Power – duration) | D (mm) | SW (mm) |
| --- | --- | --- | --- |
| Computer | $20 \pm 10$ g, 25 W–30 s | 5.86 | 8.96 |
| Ikeda *et al* [23] | 22 g, 30 W–60 s | $6.7 \pm 0.8$ | $7.02 \pm 0.6$ |
| Leshem *et al* [38] | 10 g, 30 W–30 s | $5.95 \pm 1.3$ | $8.99 \pm 2.1$ |
| Nakawaga *et al* [36] | $18.1 \pm 7.6$ g, 30 W–30 s | $5.6 \pm 1.4$ | $7.3 \pm 1.1$ |

We also found reasonable agreement between our computational results and those reported by Shah *et al* [21] in an *in vitro* model that mechanically simulated catheter displacement under conditions roughly similar to ours: perpendicular catheter, 20 W, 50 cycles/minute reaching a peak “systole” of 20 g and a nadir “diastole’ of 10 g. Shah *et al* concluded that contact force–time integral (FTI) is a predictor of lesion depth at constant RF power in a contractile bench model simulating the beating heart. In terms of FTI, our model predicts a lesion depth of 6.66 mm for FTI of 900 g·s (static case with 30 g and 30 s) and 6.08 mm for FTI of 600 g·s (static case with 20 g and 30 s, and dynamic case with 20 ± 10 g and 30 s), while Shah *et al* reported a value ∼ 6 mm for FTI of 900 g·s and ∼5.2 mm for FTI of 600 g·s. Note that although they used 60 s ablations and we simulated only 30 s, using FTI allows the comparison.

No specific experimental data are available as regards comparing the static and dynamic cases. Although Shah *et al* [21] compared the lesion sizes of static contact and variable/intermittent (dynamic) contact, both groups had really very different FTI values, with significant differences in lesion size. However, Leshem *et al* [38] did find similar lesion depths from a non-beating model based on thigh muscle and a beating heart model (both *in vivo*) in case of standard energy setting (30 W for 30 s), which indicates that the static and dynamic models might be equivalent in terms of predicting lesion depth in the case of standard energy setting.

Finally, we found that lesion depth is relatively independent of electrode insertion depth when measured from the electrode tip rather than from the tissue surface, and obviously with the electrode inserted (see D* in Fig. 3). Specifically, D* values were 4.55, 4.62 and 4.61 mm for the static case with contact forces of 10, 20 and 30 g, respectively, and 4.64 mm for the dynamic case (20 ± 10 g contact force). Interestingly, the difference between the D* values and those shown in Table 2 for the inserted electrode (i.e. D values) is due almost entirely to the insertion depths, i.e. 0.86, 1.46 and 2.05 mm for the static case with contact forces of 10, 20 and 30 g, respectively. This is reasonable because no pressure-dependence of the electrical and thermal conductivity was considered. Within the ranges considered this suggests that no matter how much force the electrode exerts on the tissue, the thermoelectric result around the electrode is practically the same, so that lesion depth is almost the same in terms of compressed tissue when measured from the electrode tip. In this regard, Ikeda *et al* [23] found by regression that lesion depth increases by 0.09 mm/g. From our data in Table 2 we can deduce that the electrode is inserted 0.06 mm/g, and therefore the lesion depth measured from the surface also increases 0.06 mm/g, a very similar result to the experimental findings in Ikeda *et al*. It should be remembered that for each gram of contact force, the increase in insertion depth is directly related to tissue elasticity, i.e. to its Young’s modulus. If we had used a lower value than E = 75 kPa we could have found increased lesion depths around the values reported by Ikeda *et al*.

### 4.3. Comparison with previous computational studies

As regards previous computational modeling studies that included mechanical deformation there is also some consistency with the data reported by Petras *et al* [15]. While our results show an increased lesion depth of 1.2 mm when CF is raised from 10 to 30 g, their *ex vivo* and computational experimental results (at 20 W−30 s) rise by 0.8 mm when CF goes from 10 to 20 g, and 1 mm when it increases from 10 to 40 g. The differences could be due to the way of quantifying lesion size, since the depth values reported by Petras *et al* (2.3−3.3 mm) seem peculiarly small compared to those obtained experimentally in other experimental studies using 20 W-30 s (∼4 mm), for example in Guerra *et al* [39]. It is not possible to make a direct comparison with the computational results of Yan *et al* [16] as they modeled a constant voltage ablation. However there is also some consistency, since they found that lesion depth increased by 0.85 mm when CF went from 10 to 30 g (variable power between 27 and 30 W for 30 s). Nor can we compare them with Singh and Melnik’s results [17] since they did not report lesion depths and their ablations lasted twice as long (60 s). Nor can we compare them with Ahn and Kim’s results [18] since they used a level of power much lower than the one used in the clinic (<10 W).

### 4.3. Limitations of the study

Since there is still no mathematical model of the behavior of the gas bubbles formed in tissue at high temperatures that cause steam pops at a temperature of 100 ºC, our model cannot predict this phenomenon. Instead, we modeled the behavior of tissue around 100 ºC using state-of-the-art mathematical RFCA modeling, i.e. a drop in electrical conductivity to model tissue desiccation, and the enthalpy method to model the energy balance associated with phase change (vaporization). Although our computer simulations did not stop when tissue temperature reached 100 ºC, these two features kept the maximum temperature below 105 ºC and predicted lesion depth reasonably well compared to previous experimental results based on a beating heart model and similar conditions [23,36,38]. Interestingly, steam pop incidence was very low in these experimental results: 13% in Ikeda *et al* [23] and none in Leshem *et al* [38] and Nakagawa *et al* [36]. Although this is somewhat speculative, we think that the specific conditions we simulated, i.e. a standard energy setting (25 W for 30 s) and moderate contact force (<30 g), are not liable to superheat and cause detectable/audible steam pops (even though a relatively large tissue area reached 100 ºC during the simulations, as shown in Fig. 4). In contrast, Ikeda *et al* [23] did report increased incidence of steam pops at a higher power (40 W) and greater contact force (50−100 g). Our conclusions are therefore only valid for conditions not prone to overheating.

Second, only a perpendicular catheter was considered, which allowed axial symmetry and simplified the problem to a 2D model. Third, the thermal effect of the circulating blood was modeled with convection transfer coefficients instead of solving the velocity map in the blood. This simplification is suitable for predicting lesion depth and maximum tissue temperature, and although it tends to overestimate the surface width [13], it is suitable in the context of comparing static vs. dynamic insertion depths. Fourth, electrode irrigation was modeled assuming constant electrode temperature. It is important to point out that this approach may not be suitable for cases in which the CF is high or the irrigation holes are partially blocked by tissue [8]. As already mentioned, our conclusions should therefore be considered valid only at low contact forces and unhampered irrigation.

Fifth, only contact force variations were associated with heartbeats since they are more relevant than others such as breathing. Only a frequency of 60 beats per minute (normal rhythm) and a sinusoidal waveform were simulated. This waveform was chosen since it is easy to synthesize mathematically for application as a time-changing boundary condition. Although the actual evolution of CF shows a waveform a bit more pointed than a simple sinusoidal [40], we think that there is quite a reasonable similarity between both waveforms and that the choice of a sinusoidal is a good approximation that should not invalidate the conclusions. And sixth, viscous behavior was not included in the mechanical model.

## 5. Conclusions

We developed an RFCA dynamic model including heartbeat-induced electrode displacement at a standard energy setting and moderate contact force. The model predicts lesion depth reasonably well as compared to previous experimental results based on a beating heart model. Under these specific conditions, the lesion size computed by a dynamic model was practically identical to that of a static model in which the electrode remained constantly inserted at an average depth.

## Appendix. Thermal characteristics for dehydrated tissue

Once the temperature reached 100 °C, we assumed that the tissue lost its aqueous component and became dehydrated. The specific heat, thermal conductivity and density of dehydrated tissue are shown in Table 1. These were estimated as described in this Appendix. In all that follows, the subscript *w* refers to the properties of water, *dh* to dehydrated tissue, and the absence of subscript to hydrated tissue (whose characteristics are shown in the Table 1). Let us assume that in 1 kg of hydrated tissue we have *C* kg of water and (1−*C*) kg of dehydrated tissue, so that the volume (*V*) of each of the parts is:

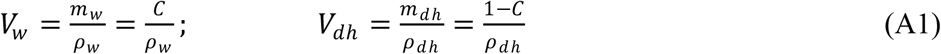

The volume of hydrated tissue will be the sum:

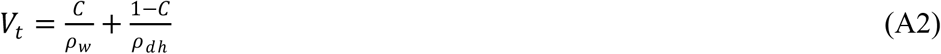

Consequently, tissue density will be:

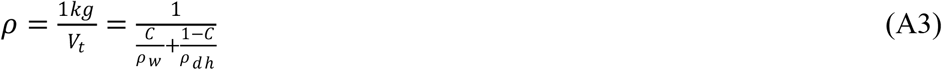

and density of hydrated tissue will be:

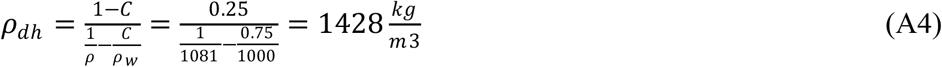

Since the specific heat measures the energy needed to raise a 1 ºC of matter, if we have 1 kg of (hydrated) tissue, *C* kg is water and (1−*C*) kg is dehydrated tissue. If we increase the tissue temperature by 1º C we can separate the energy needed for each component (water and dehydrated tissue) as follows:

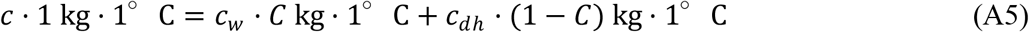

where we can obtain the specific heat of dehydrated tissue:

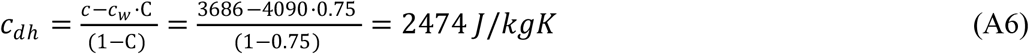

The thermal conductivity of dehydrated tissue was estimated using the models described by Carson [30] for the equations of non-frozen and non-porous food (since *k*_*w*_/*k*_*dh*_ was ≈3, as demonstrated below). Although Carson describes several models for this particular case, he also recognizes that similar results are obtained regardless of the one used. For this reason, we used the ‘parallel model’, where *k* can be expressed as:

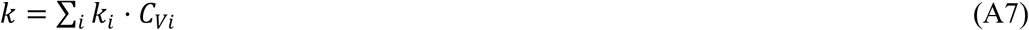

where *k*_*i*_ is the thermal conductivity of each constituent and *C*_*Vi*_ the volumetric ratio in the mixture. Firstly we must calculate the volumetric ratio of water from the mass fraction (*C*):

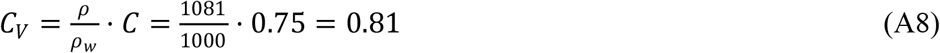

and then apply the parallel model

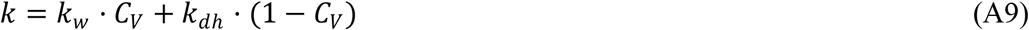

where *k*_*w*_ = 0.624 W/m·K [31], to finally obtain the thermal conductivity of dehydrated tissue (data assessed at 37 ºC)

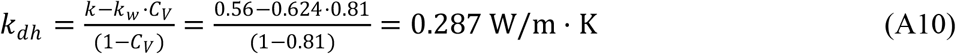

which provides a relation *k*_*w*_/*k*_*dh*_ = 2.17 and suggests that the chosen model is suitable because it is close to 3. This decrease in thermal conductivity once tissue is completely dehydrated is qualitatively consistent with reports from experimental studies using liver (from 0.5 to 0.19 W/m·K) [32] and swine left ventricle samples (from 0.61 to 0.50 W/m·K) [33] as used in previous numerical studies [25,32].

